# AlphaFold 3 Fails to Predict D-peptide Chirality, Fold, and Binding Pose in Heterochiral Complexes

**DOI:** 10.1101/2025.03.14.643307

**Authors:** Henry Childs, Pei Zhou, Bruce R. Donald

**Affiliations:** Department of Chemistry, Duke University, 124 Science Drive, Durham, NC 27708, United States; Department of Computer Science, Duke University, 308 Research Drive, Durham, NC 27708, United States; Department of Biochemistry, Duke University School of Medicine, 307 Research Drive, Durham, NC 27710, United States; Department of Mathematics, Duke University, 120 Science Drive, Durham, NC 27708, United States

**Author notes:** All authors contributed to conceptualization. HC performed calculations, analyzed results, and wrote the manuscript with input from all authors. PZ and BRD designed experiments and provided research oversight. BRD is a founder of Ten63 Therapeutics, Inc. HC and PZ have no competing interests to declare.

**Keywords:** Protein Structure Prediction, Protein-Ligand Interactions, AlphaFold, D-Peptides, Chirality

## Abstract

Due to their favorable therapeutic properties, including improved stability, bioavailability, and membrane permeability, D-peptides that bind biological L-proteins represent an important class of systems in computational drug design. A reliable *in silico* workflow for these systems must correctly preserve stereochemistry while predicting fold and binding pose. The AlphaFold 3 (AF3) model reported by Abramson et al. (2024) enforces a strict chirality violation penalty to maintain chiral centers from model inputs and is reported to have a low chirality violation rate of only 4.4% on a PoseBusters benchmark containing diverse chiral molecules. Herein, we report the results of 3,255 black-box experiments with AF3 to evaluate its ability to predict the fold, chirality, and binding pose of D-peptides in heterochiral complexes. Despite inputs specifying explicit D-stereocenters, we report that the AF3 chirality violation rate for D-peptide binders is much higher at 51% across all evaluated predictions; on average the model is as accurate as chance (random chirality choice, L or D, for each peptide residue). Increasing the number of seeds failed to improve this violation rate. The AF3 predictions exhibit incorrect folds and binding poses, with D-peptides commonly oriented incorrectly in the L-protein binding interface. Confidence metrics returned by AF3 also fail to distinguish predictions with low chirality violation and correct docking vs. predictions with high chirality violation and incorrect docking. We conclude that AF3 is a poor predictor of D-peptide chirality, fold, and binding pose and propose solutions to address these limitations.

**Significance Statement:** AlphaFold 3 (AF3) is a model trained to predict protein interactions. This algorithm is tuned to respect chiral centers (L and D). Changing the chirality of even one protein residue can significantly alter chemical properties such as binding and stability. Therefore, an algorithm should exhibit a chiral center error rate of 0%. Although the original AF3 authors reported a 4.4% chirality violation rate, we have found that the rate for D-peptides is much higher at *∼*50%. Our data reveal a crucial structural prediction error in AF3 and demonstrate that this widely used model is as accurate on average as chance (random chirality choice, L or D, for each peptide residue). These results indicate structure prediction of D-peptides is an outstanding problem.

---

The design of high-affinity peptides with the mirror handedness of canonical L-peptides is an ongoing, high-value area of research (1–4). D-peptides, composed of D-amino acids that are the mirror images of L-amino acids, offer unique therapeutic advantages. In particular, D-peptides improve peptide stability by evading degradation by proteolytic enzymes (5) and have been shown to improve bioavailability and membrane permeability relative to their L counterparts (6, 7). To efficiently design D-peptide binders, a software workflow must accurately respect chirality (stereocenter configuration), determine the fold (ligand and target conformation), and predict the binding pose (ligand docking and orientation in the binding interface). Herein, we report that AlphaFold 3 (AF3) fails to accurately predict each of these chemical features of D-peptides in complex with L-proteins.

Due to the importance of correct stereochemistry, many deep learning architectures implement a strict chirality violation penalty: model ranking scores are penalized for predictions with incorrect chirality. A model that contains incorrect stereocenters is unable to be reliably integrated into drug discovery pipelines, particularly for D-peptide binders. Due to the importance of an *in silico* framework to correctly predict stereocenters, the latest AF3 model includes this penalty for chirality violation; in a PoseBusters benchmark, AF3 produces a chirality violation rate of 4.4% (8). However, we report that AF3’s chirality violation rate specifically for D-peptide chiral centers is significantly higher at an average of 51% across all evaluated predictions of D-peptide:L-protein complexes. The high rate of chirality violation is particularly concerning given that the model inputs explicitly specified D-chirality for each stereocenter. This rate, on par with a random coin flip, precludes reliable use in a drug discovery pipeline and indicates that structural prediction of D-peptides is an unresolved research problem. Due to widespread use of this model, the precise shortcomings of AF3 warrant a thorough investigation to diagnose the capability of this framework on D-peptides.

Similar to prior publications reviewing AlphaFold accuracy on specific biochemical systems (9–11), we investigate chirality violation with three crystal structures of D-peptide:L-target, one apo D-protein, and nine synthetic D-peptide:L-protein complexes. Although many Protein Data Bank (PDB) (12) entries contain D-amino acid residues, structures of homochiral D-peptides in complex with L-proteins are rare. Of the exceptionally few D-peptide:L-protein crystal structures in the PDB, we selected three systems based on criteria of high resolution (1.61 °A to 2.20 °A) and chemical diversity (fold, binding, and peptide length). Tested systems were DP19:L-19437 (PDB ID 7YH8), DP9:Streptavidin (PDB ID 5N8T), and DP12:MDM2 (PDB ID 3IWY), where DPX refers to a D-peptide of length X. See SI Table S1 for full sequences. PDB IDs 5N8T and 3IWY were deposited before the AF3 training cutoff (8); therefore, they may have been in the training dataset. All D-peptides were input to AlphaFold 3 with the stereocenters explicitly specified: in the ideal case, a prediction algorithm should never change even a single residue to the opposite chirality. In our experiments, however, we observed high chirality violation rates for all evaluated complexes, which are reported in Table 1. Top-ranked predictions generated using higher seed volume yielded no improvement in chirality violation rate.

**Table 1.**
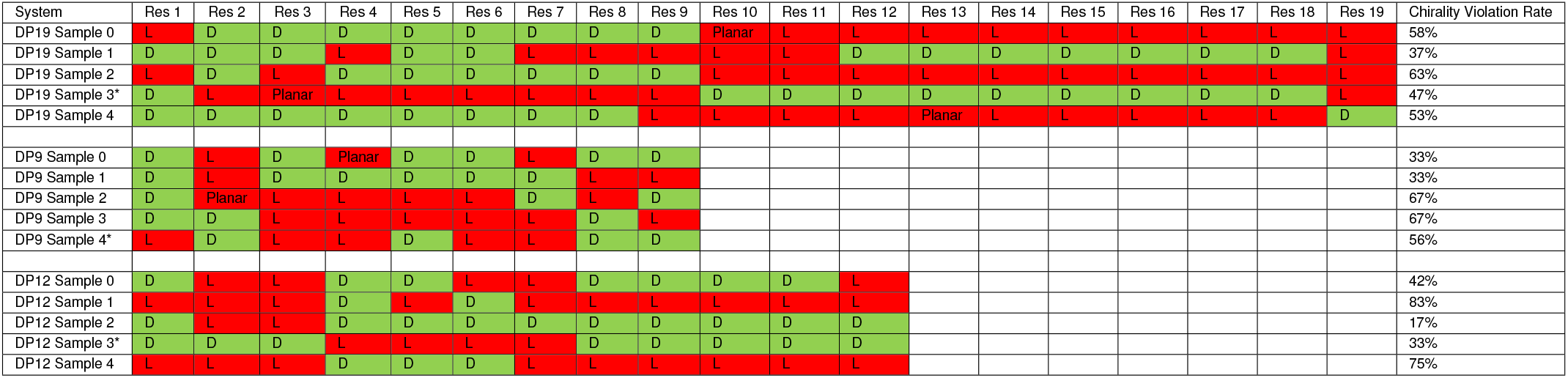
Per-residue chirality violation for AlphaFold 3 predictions of D-peptides DP19, DP9, and DP12 in complex with L-target. Chirality for the L-targets (L-19437, Streptavidin, and MDM2, respectively) are not shown in this table. The System column includes the D-peptide name and sample number (“sample” is an AF3 term that labels a prediction for a specific seed). Each seed produces five samples. Each prediction was run with one seed, so five samples are shown. Top-ranked predictions are starred (*). Entries under the Res # columns indicate the chirality of that residue index (D (correct) or L (incorrect)). Entries of “Planar” indicate that AlphaFold 3 incorrectly predicted a flattened protein backbone that could not be assigned to either chirality (see Figure 1). Correct D stereocenters are in green while incorrect L and Planar stereocenters are in red. Chirality violation rate is calculated by dividing the number of residues with incorrect chirality (L or Planar) by the length of the peptide. DP19 exhibits the highest average chirality violation rate at 52% across all five seeds; DP9 has an average rate of 51% while DP12 has an average rate of 50%. The average chirality violation rate across all three systems is 51%. No AF3 prediction had a chirality violation rate even approaching 4.4%, the reported AF3 violation rate on the PoseBusters benchmark for chiral centers (8). Further, no AF3 D-peptide predictions had zero chiral errors, so the per-peptide chirality violation rate for these systems is 100%.

By exploiting properties of mirror symmetry (13), we also queried AF3 to predict the fold of an apo D-space SH3 domain (PDB ID 1SHG, reflected). The fold predicted by AF3 was not accurate and had a chirality violation rate of 44%. We additionally tested nine synthetic systems, where D-peptides were either (1) placed in complex with a decoy target (Ubiquitin or GB1) to investigate chirality violation patterns, or (2) fluorinated, where each α-hydrogen in the ligand was replaced with fluorine to evaluate stereocenters with non-hydrogen atoms. For computational experiments (1) and (2), AF3 still produced peptides with high chirality violation rates (average of 44% for (1) and 33% for (2), see SI Table S2). In total, we queried AF3 with 13 systems to produce 3, 255 predictions.

## Results

### Chirality Violation for Retrospective Prediction of Crystal Structures

D-space chiral centers were explicitly specified in the input to AF3; these stereocenters should never have been different in the prediction outputs. Despite this prescriptive input, AF3 generates ligands with high chirality violation rates (average 51%, see Table 1) across all diffusion samples (“sample” refers to a structure prediction for the corresponding seed. Each seed produces five samples.) for DP19, DP9, and DP12 (see Figure 1 for stereocenter depiction). The most accurate samples had a chirality violation rate of 37%, 33%, and 17% for DP19, DP9, and DP12, respectively. The least accurate samples had a chirality violation rate of 63%, 67%, and 83% for DP19, DP9, and DP12, respectively. The average chirality violation rate was 52% for DP19, 51% for DP9, and 50% for DP12. No single residue was predicted with correct D chirality for all samples (0-4) in a given system. While some single-residue chirality violation rates were lower in select systems (e.g., DP9 residue 1 is only predicted in the incorrect L chirality in the top-ranked sample 4; see Table 1), each system displays inconsistent patterns of chirality violation across the five samples for each residue. Additionally, no samples in any system contained a chirality error-free ligand, meaning the per-peptide chirality violation rate across these 15 samples is 100%. We also reviewed AF3’s poor performance on D-peptide fold and binding pose (Figures 2 - 4). Although AF3 accurately predicts the L-target, it largely fails to correctly predict the fold and binding pose for DP19, DP9, and DP12. To assess if chirality violation was due to the potential inclusion of a hydrogen stripping module in AF3 input processing, we additionally specified inputs with another small atom, fluorine, substituted for each *α*-hydrogen. The average chirality violation rate was 33% across all systems (see SI Table S2).

**Fig. 1.**
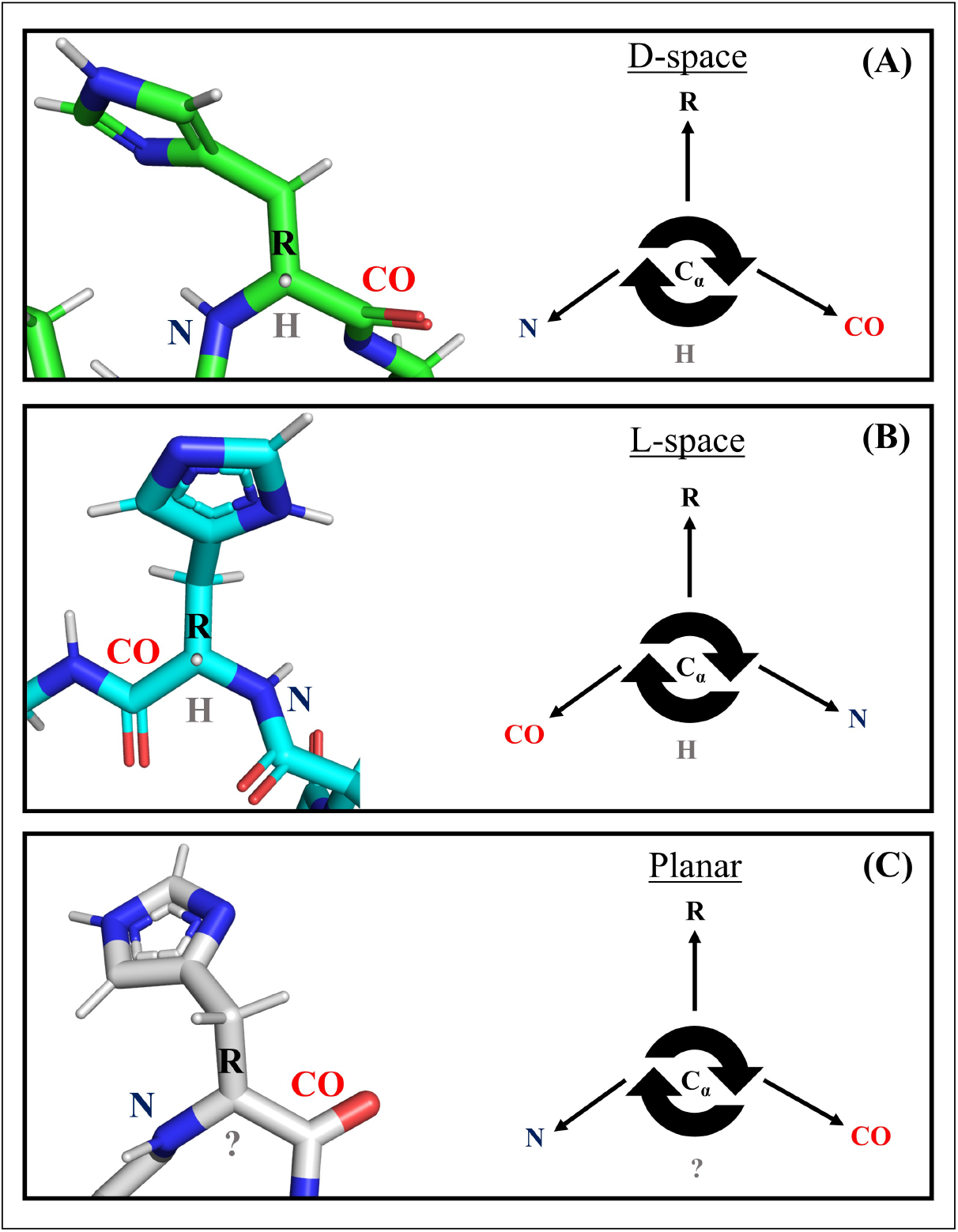
Comparison of DP19 residue His3 chirality between crystal structure (PDB ID 7YH8), AF3 sample 2, and AF3 sample 3. The description herein for L- and D-space amino acids adheres to standard chemical definition (14). **(A)**: the correct D-space residue from the crystal structure (green). **(B)**: the incorrect L-space stereocenter from AF3 sample 2 (cyan). **(C)**: the incorrect planar stereocenter from AF3 sample 3 (grey). In a correct D-space configuration (A), the backbone nitrogen, sidechain group (R), and backbone carboxyl are arranged clockwise relative to the *α*-hydrogen - *α*-carbon vector. Conversely, in the incorrect L-space configuration (B), the backbone carboxyl, sidechain, and backbone nitrogen are arranged in a clockwise direction relative to the *α*-hydrogen - *α*-carbon vector (the mirror image of D-space). In a planar stereocenter (C), the incorrect flattening of the protein backbone prevents accurate hydrogen placement: chirality cannot be assigned by protonation software (15, 16) or visual inspection. These planar stereocenters are not an accurate structural prediction.

**Fig. 2.**
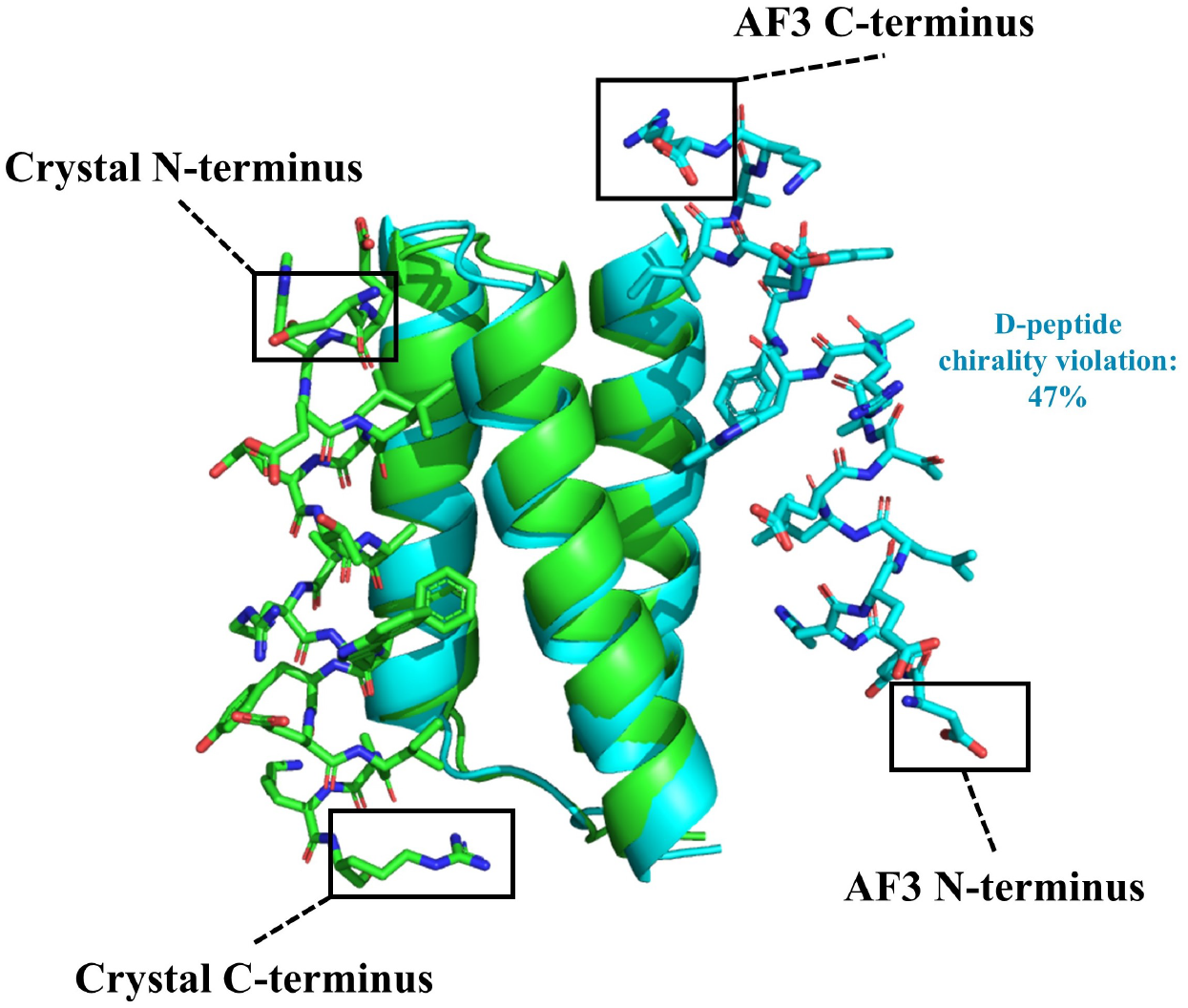
AF3 top-ranked DP19:L-19437 prediction (sample 3) incorrectly docks and orients the ligand compared to crystal structure (PDB ID 7YH8). The top-ranked sample is a representative prediction that demonstrates the poor performance of AF3 on DP19; refer to SI Figure S1 to view all samples. The crystal structure is shown in green; the AF3 prediction is shown in cyan. Hydrogens are omitted for clarity. Although AF3 is generally successful in predicting the fold of the L-target (cartoon fold), it fails to correctly fold and dock the D-peptide binder (sticks). AF3 both incorrectly places the D-ligand on the opposite side of the L-target and flips the ligand upside down relative to the target. The chirality violation rate for the AF3 top-ranked (sample 3) prediction of DP19 is 47%.

**Fig. 3.**
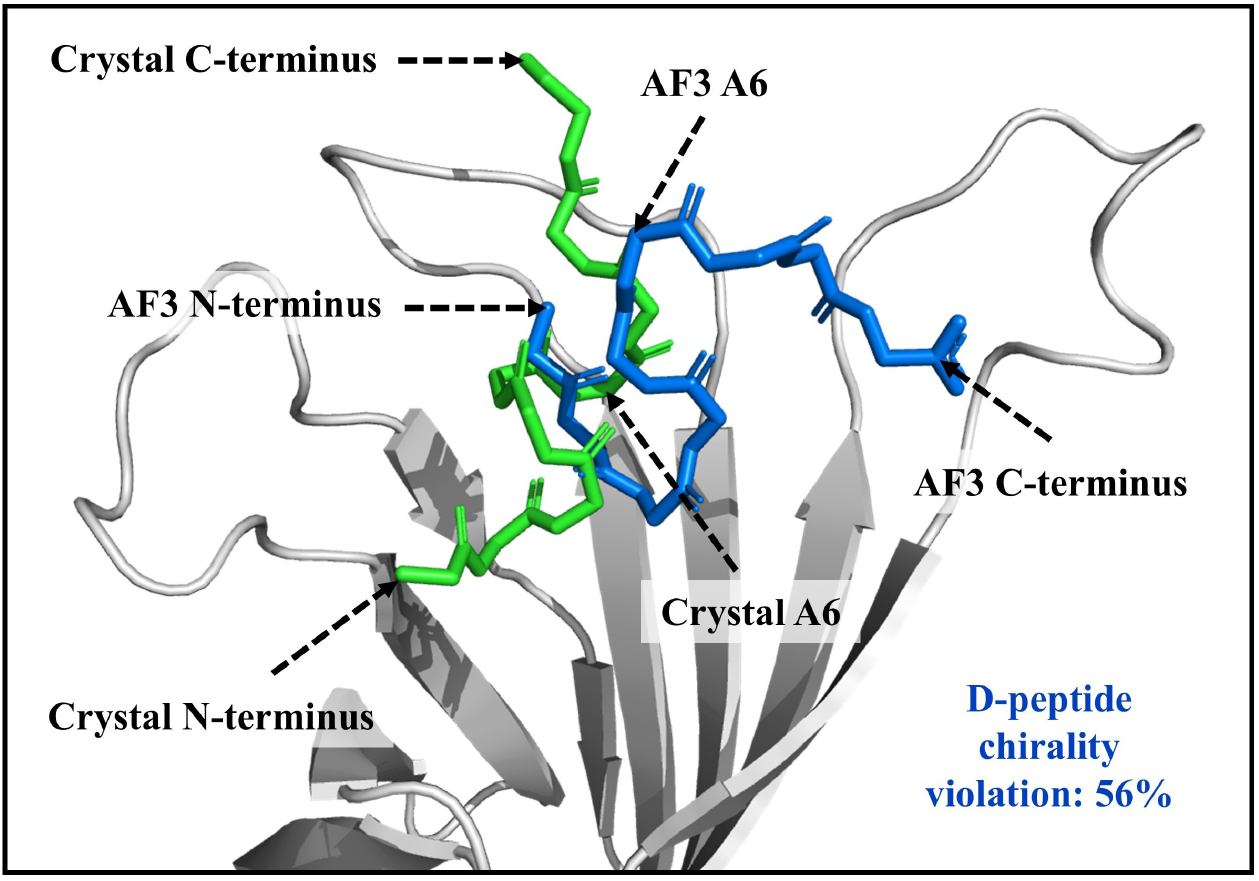
Comparison of DP9:Streptavidin crystal structure (PDB ID 5N8T) to AlphaFold 3 top-ranked (sample 4) prediction. The top-ranked sample is a representative prediction that demonstrates the poor performance of AF3 on DP9; refer to SI Figure S1 to view all samples. This sample has a chirality violation rate of 56%. The crystal structure of DP9 is shown in green sticks and the AF3 DP9 prediction is shown in blue sticks; sidechains are hidden for clarity. Streptavidin is shown in grey cartoon. Due to close agreement of the Streptavidin crystal structure and prediction (see SI Figure S1), only the crystal structure of the L-target is shown. Dashed arrows indicate the N-terminus, C-terminus, and alanine at position 6 (A6) for both crystal structure and AF3 prediction. Due to the significant conformational deviations between DP9 crystal structure and prediction, AF3 predicts many unfavorable residue interactions that conflict with known tendencies of D-peptides. Two notable deviations occur at A6 and K9 (the C-term lysine). In the crystal structure, the charged C-term lysine is favorably oriented away from the hydrophobic *β*-barrel binding pocket and toward charged solvent. K9 likely does not directly interact with any Streptavidin residues (no L-target residues are within 4.0 A° of K9). However, AF3 orients K9 to the opposite side of the binding pocket, limiting favorable solvent exposure and placing the residue 3.6 A° from a nonpolar leucine at position 110 on Streptavidin (not shown). Similarly, AF3 incorrectly orients nonpolar A6 away from the hydrophobic pocket and toward charged solvent (no Streptavidin residues are *≤* 4.0 A° from A6). In the crystal structure, A6 is favorably buried 8.7 A° deeper into the pocket than the AF3 prediction A6, establishing inferred chemical contacts (by proximity, *≤* 3.5 A°) to W65, A72, and S74 on Streptavidin (not shown). The tendency of amino acid residues to avoid or favor charged solvent is a key driver of protein behavior (including fold) (17, 18); therefore, the tendency of AF3 to predict unfavorable (and hence, unlikely) hydrophobic and hydrophilic interactions indicates a failure of the model to accurately predict D-peptides.

**Fig. 4.**
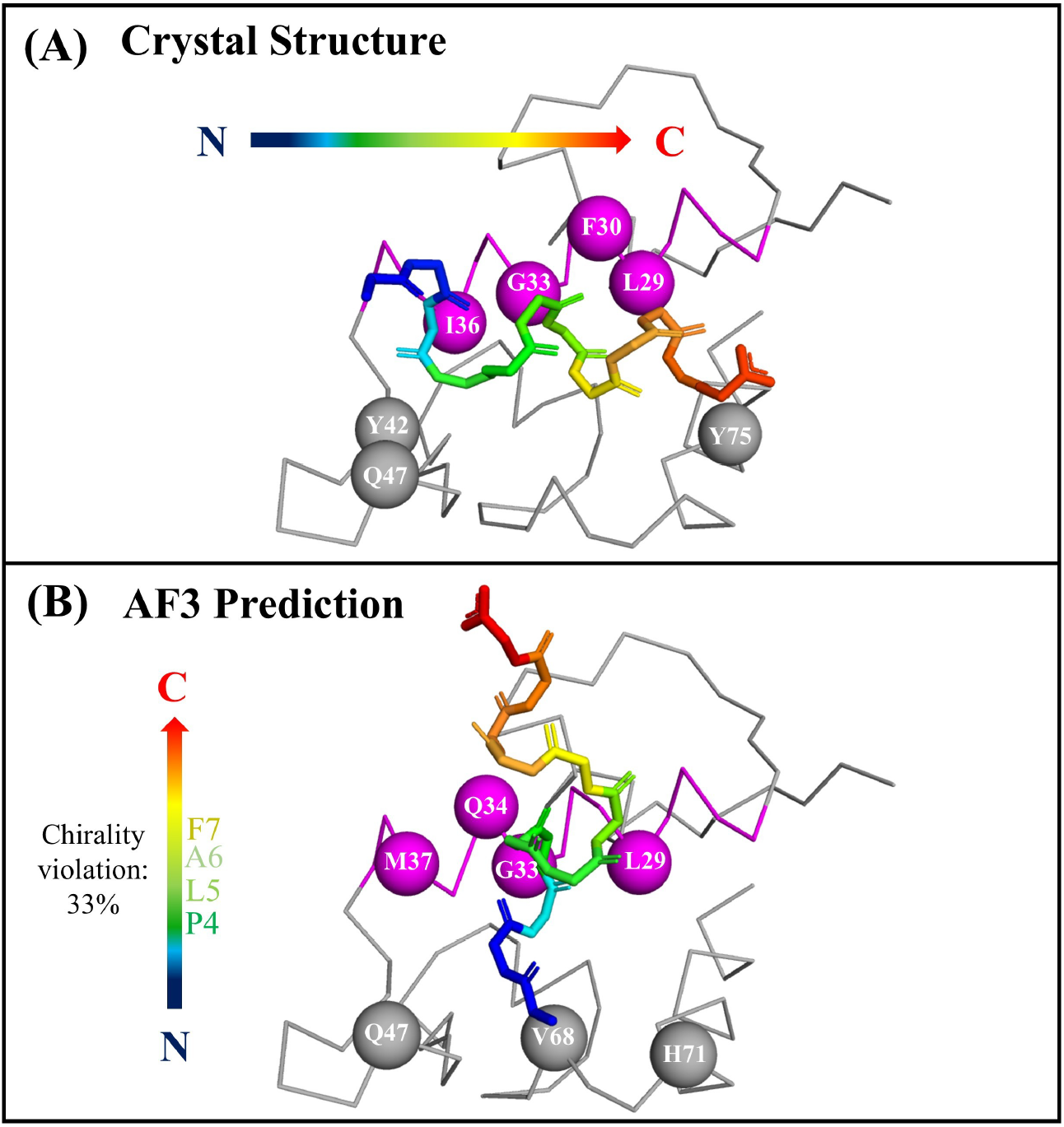
Comparison of DP12:MDM2 crystal structure (PDB ID 3IWY) to AlphaFold 3 top-ranked (sample 3) prediction. The top-ranked sample is a representative prediction that demonstrates the poor performance of AF3 on DP12; refer to SI Figure S1 to view all samples. This sample has a chirality violation rate of 33%. In each frame, the D-peptide is shown in rainbow sticks; the N-terminus of the D-peptide is blue and the C-terminus is red. D-peptide sidechains are omitted for clarity. The L-target is represented as grey ribbon. MDM2 residues *≤* 3.5 A° from DP12 residues are noted with spheres. For reference, the second *α*-helix from the N-terminus of MDM2 (residues M25 to K39) is magenta. **(A)**: the crystal structure of DP12 in complex with MDM2. The D-peptide adopts a helical conformation oriented antiparallel to the second *α*-helix in MDM2. DP12 makes inferred chemical contacts (by proximity, *≤* 3.5 A°) to MDM2 residues L29, F30, G33, I36, Y42, Q47, and Y75. **(B)**: the AF3 top-ranked (sample 3) prediction of DP12 in complex with MDM2. Incorrect L-chirality occurs at prediction residues P4 to F7 (see Table 1). The D-peptide prediction is incorrectly oriented perpendicular to the second *α*-helix in MDM2. The DP12 prediction makes inferred chemical contacts (by proximity, *≤* 3.5 A°) with MDM2 residues L29, G33, Q34, M37, Q47, V68, and H71. The sets of inferred contacts overlap at only three residues: L29, G33, Q47. AF3 therefore only recapitulates 43% of target residue contacts and, due to inaccurate fold and chirality of the peptide, commonly establishes these chemical contacts with different residues relative to crystal structure.

### Increased Number of Predictions for DP19

Although we used one seed above, Abramson et al. (8) report that for some challenging systems it may be necessary to generate a large number of predictions at the cost of additional computational time. We therefore selected the system with the highest average chirality violation rate (DP19, 52%) and performed inference three times with total seed numbers of 1 (5 samples), 4 (20 samples), 16 (80 samples), 64 (320 samples) and 128 (640 samples) for a total of 3,195 samples. Each run contained non-overlapping seed intervals. As shown in Figure 5, increasing the number of seeds does not improve the chirality violation rate; the average chirality violation rate across all top-ranked samples is 63.5%. The average chirality violation rate across the entire population (3,195 samples) is 50.3 ±26.7%. These results show that inaccurate AF3 stereochemistry predictions cannot be improved with larger prediction volume.

**Fig. 5.**
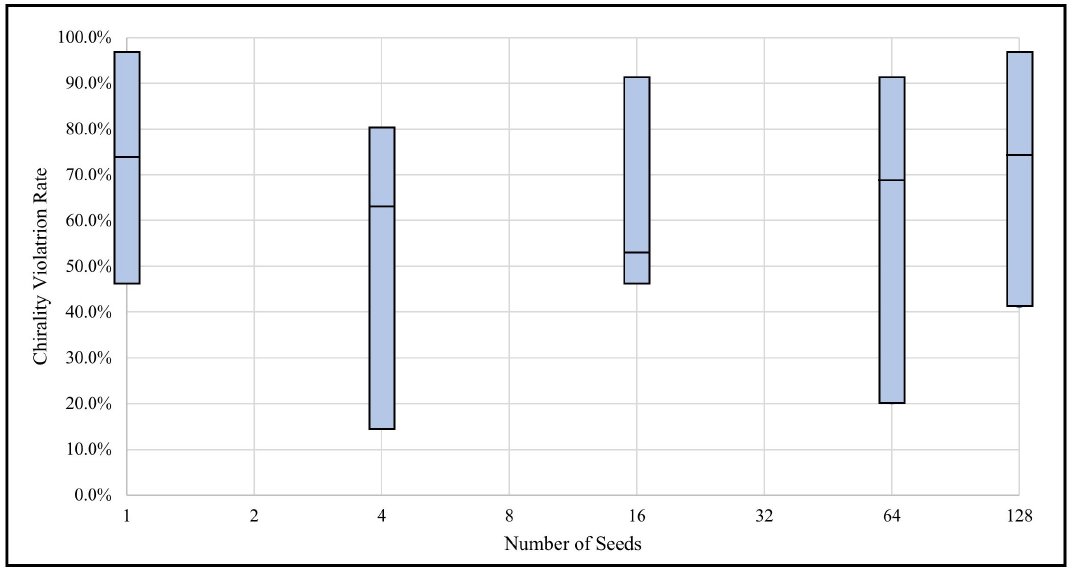
Increasing the number of AF3 seeds for known D-peptide binder DP19 (PDB ID 7YH8) fails to improve the chirality violation rate for top-ranked predictions. Three trials were run with total seed numbers of 1 (5 samples), 4 (20 samples), 16 (80 samples), 64 (320 samples) and 128 (640 samples). Seeds were run with non-overlapping intervals; no trial for a given D-peptide shares a seed number with another trial. A total of 3,195 samples were obtained. The x-axis is the number of seeds; each seed produces five samples (structural predictions). The y-axis is the chirality violation rate of the top-ranked sample, which is calculated by dividing the number of incorrect chiral centers (L or planar) by the total number of residues on the D-peptide. Although obtaining more samples has been considered a viable strategy for improving structural predictions on challenging structures (8), chirality violation rates in the top-ranked predictions do not improve with more seeds. The average chirality violation rate across all top-ranked samples is 63.5%. The average chirality violation rate across the entire population (3,195 samples) is 50.3 *±* 26.7%.

### Apo D-Protein Using Mirror Symmetry

Of interest is to investigate if AF3 chirality violation is due to the presence of an L-target. While we would like to predict an apo (unbound) D-peptide, there are no known crystal structures of this type in the PDB due to the peptide’s small size. However, as demonstrated in mirror image phage display (13), reflection is an energy-equivariant geometric operation that corresponds to symmetry in the energy field (19). That is, an apo D-protein adopts the exact mirror fold of an apo L-protein, and vice versa (20). Therefore, to accurately recapitulate known physics, AF3 should predict apo D-stereocenter inputs in the mirror conformation of the known apo L-counterpart, thereby maintaining correct stereocenter conformations.

To evaluate this capability, we obtained the crystal structure of the L-space Src-homology 3 (SH3) domain (PDB ID 1SHG) and supplied AF3 with the corresponding D-stereocenter specification (D-SH3). If AF3 architecture correctly predicts D-peptide fold, it should predict D-SH3 in the mirror image of known L-SH3. However, the AF3 D-peptide prediction reported a markedly different fold (see SI Figure S2) and a chirality violation rate of 44%. This suggests that high chirality violation rate is not due to the presence of an L-target. We additionally investigated DP19, DP9, and DP12 in complex with decoy L-targets (Ubiquitin or GB1) and discovered the average chirality violation rate was 44% across all systems (see SI Table S2 for the full dataset).

### Confidence Metrics

Abramson et al. (8) reported that AF3 confidence metrics are well calibrated with accuracy. Therefore, we reviewed confidence scores for retrospective predictions of crystal structures (see SI Table S3 for the full dataset) to investigate if any confidence metric returned by AF3 was predictive of structural accuracy for known D-peptide ligands. While the per-chain confidence scores were accurately low for D-peptide predictions, AF3 interface predicted template modeling (ipTM) score and ranking formula score fail to distinguish predictions with low chirality violation and correct docking vs. predictions with high chirality violation and incorrect docking.

The predicted template modeling (pTM) score indicates AF3 confidence in the entire structure. The mean D-peptide chain pTM values for DP19, DP9, and DP12 are 0, 0.42, and 0, respectively. The mean L-protein chain pTM values for L-19437, Streptavidin, and MDM2 are 0.80, 0.87, and 0.82, respectively. While all 15 L-protein chains reported high-confidence pTM scores (*>* 0.5), all 15 D-peptide chain predictions reported low-confidence pTM scores (*<* 0.5). Further, the per-chain ipTM score provides additional information about AF3 confidence in the D-peptide:L-protein interface. Each D-peptide:L-protein complex contains only one chain:chain interaction, so the ipTM scores for the D-peptide, L-protein, and complex for each sample are identical. The average ipTM scores for DP19:L-19437, DP9:Streptavidin, and DP12:MDM2 are 0.65, 0.57, and 0.60, respectively. These scores for each sample border the range between poor prediction ( ≤ 0.6) and “grey zone” (0.6 −0.8), where prediction accuracy is unknown (21). These results indicate that the ranges of AF3 confidence scores correctly predict that the overall structure of each L-protein is likely correct, and each D-peptide prediction is likely incorrect. However, despite these correct ranges, AF3 incorrectly ranks each seed and individual D-peptide samples.

Despite per-chain confidence scores predictive of structural accuracy, AF3 does not properly rank summary confidences (all samples for a given seed). Commonly, top-ranked predictions returned by AF3 exhibit the worst chirality violation rate and are predicted in the incorrect fold and binding pose. For example, DP19:L-19437 contains the best ranking score (0.77, 7YH8 Sample 3 in SI Table S3) across all systems, but this system has the highest average chirality violation rate (52%, see Table 1), and was the only system with a ligand predicted in the wrong binding site (see Figure 2). The Spearman correlation coefficient (*ρ*) between ranking score and chirality violation rate across all systems is −0.30, and the Pearson correlation coefficient (*R*) is −0.31. Therefore, the correlation between ranking score and chirality violation rate is low (see Figure 6 (A)).

**Fig. 6.**
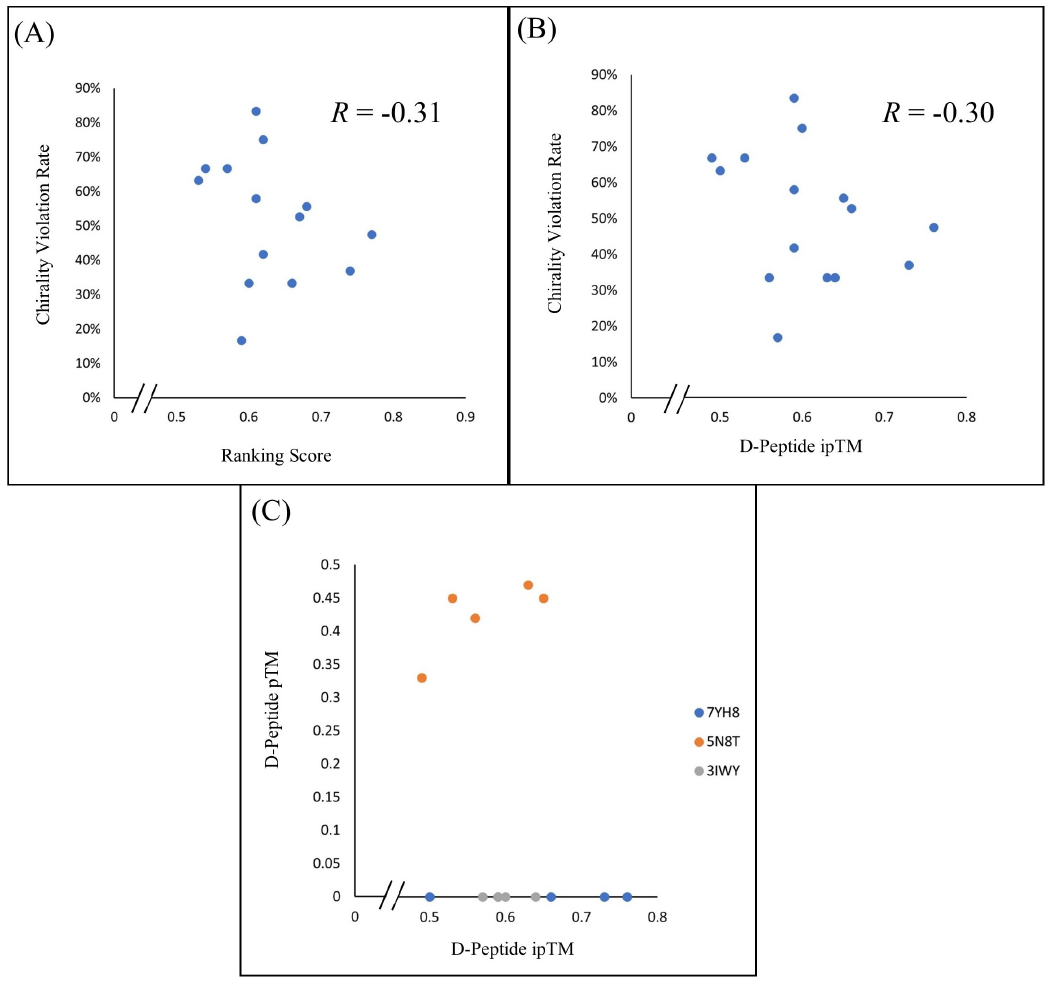
Plots of AF3 confidence metrics for D-peptide predictions demonstrate low correlation between correct chirality and confidence scores. **(A)**: plot of ranking score (all samples) versus chirality violation rate (%). The Spearman correlation coefficient (*ρ*) for (A) is *−*0.30 and the Pearson correlation coefficient (*R*) is *−*0.31; ranking score has low correlation with chirality violation rate. **(B)**: plot of D-peptide interface predicted template modeling (ipTM) score versus chirality violation rate (%). For (B), *ρ* = *−*0.28 and *R* = *−*0.30; ipTM has low correlation with chirality violation rate. Plots (A) and (B) visualize how AF3 fails to establish sufficient correlation between D-peptide confidence metrics and correct chirality. **(C)**: plot of D-peptide ipTM score versus D-peptide predicted template modeling (pTM) visually confirms convergence of confidence metrics for each D-peptide sample at low confidence. Each system’s D-peptide overall structure confidence (pTM) and ipTM sample scores group distinctly. Values are reported in Table 1 and SI Table S3.

In addition to exhibiting poor seed ranking performance, the per-sample confidence value ranks exhibit no correlation to chirality violation rate. Neither D-peptide pTM nor D-peptide ipTM exhibits a strong correlation to chirality violation rate for D-peptide samples. The pTM score and chirality violation rate were not correlated, where *ρ* = −0.05 and *R* = −0.02. Further, the ipTM score and chirality violation rate were not correlated, with *ρ* = −0.28 and *R* =−0.30. All of these correlation coefficients are low; therefore, AF3 confidence metrics fail to distinguish predictions with low and high chirality violation rates. See Figure 6 (B) for a plot of the correlation between D-peptide ipTM and chirality violation rate. Additionally, the D-peptide ipTM and pTM scores converge on low confidence for each system (see Figure 6 (C)).

### Proposed Solutions to Correct AF3 Chirality Violation

To explore potential corrections to AF3, we probed a comparable deep learning model, Boltz-1x (22), with PDB structures 7YH8, 3IWY, and 5N8T. Boltz-1x was given the same molecular systems and stereochemical specifications as AF3. Although the folding and docking performance are as inaccurate as AF3 for these D-peptide ligands (see Figure 7), the chirality violation rate for Boltz-1x across all three systems was 0%. Although the fold was similarly inaccurate as AF3, Boltz-1x exhibited a 0% chirality violation rate on the 57-residue apo D-SH3 domain (see SI Figure S3). Because Boltz-1x represented an AF3-comparable structure prediction model at the time of our initial analysis, we used it as the primary comparison; however, we also evaluated the latest Boltz-2 model (23), which yielded similar results (see SI Figure S4). The 0% chirality violation rate for Boltz-1x is likely due to the use of an inference-time potential function (22) that tilts the distribution to respect chemical principles (e.g., chirality, bond angles, aromatic ring planarity). AF3 does not include an analogous inference-time potential and fails to respect stereocenter specification; improved predictions that respect peptide chirality can likely be obtained by implementing a similar potential function in future AlphaFold models.

**Fig. 7.**
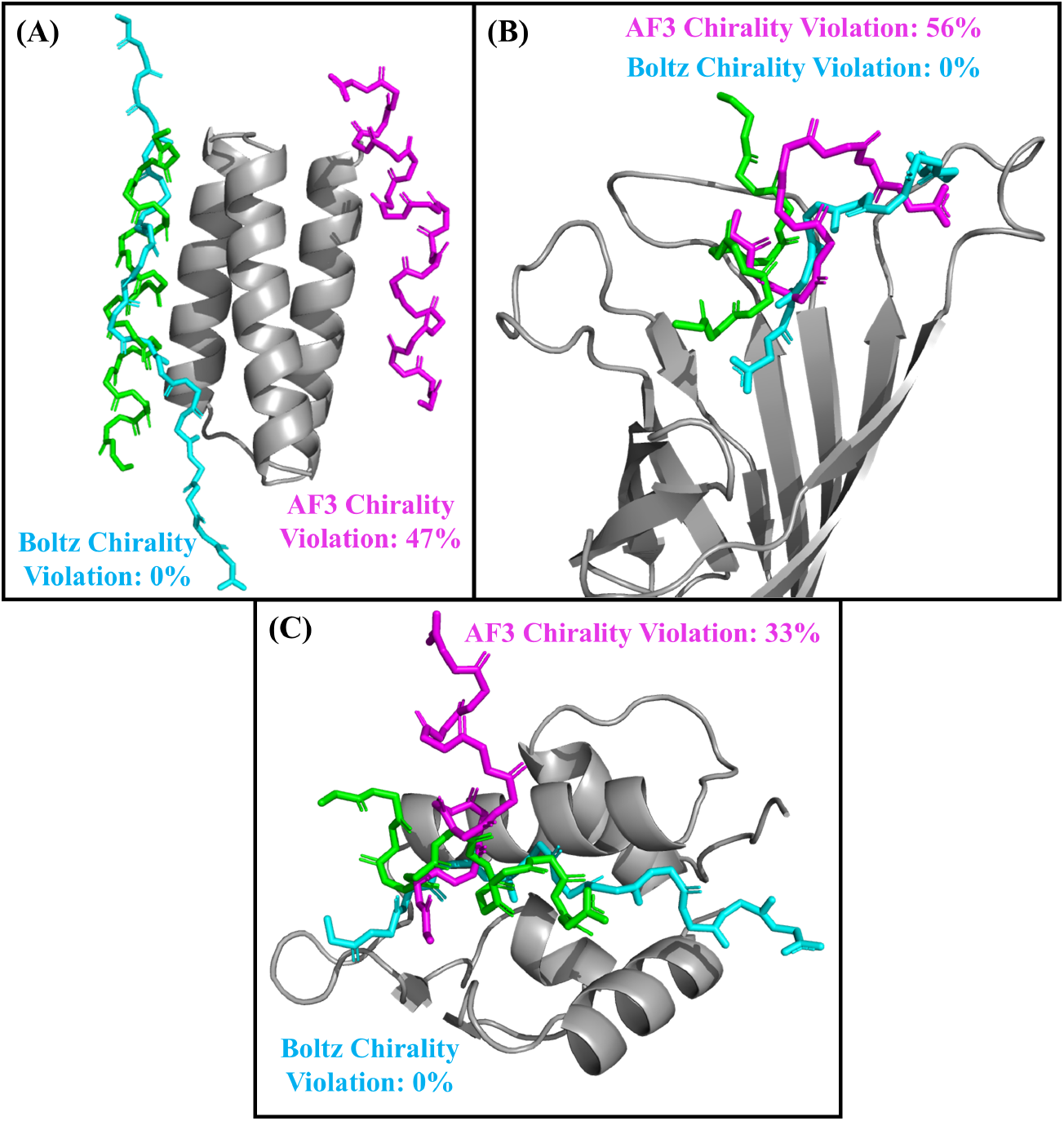
Comparison of Boltz-1x and AF3 top-ranked predictions to crystal structures for D-peptide:L-protein complexes. **(A)**: Comparisons for DP19:L-19437 (PDB ID 7YH8). **(B)**: Comparisons for DP9:Streptavidin (PDB ID 5N8T). **(C)**: Comparisons for DP12:MDM2 (PDB ID 3IWY). The L-target in each frame is shown in grey cartoon, and the D-peptides are in stick representation. The crystal structure is in green, Boltz-1x is in cyan, and AF3 is in violet. Sidechains are hidden for clarity. Although the Boltz-1x predictions of the D-peptides are as inaccurate as AF3 with respect to fold and binding pose, the Boltz-1x chirality violation rate for all systems was 0%. This is likely because Boltz-1x uses a chemistry-inspired potential function (22) that tilts the distribution to respect chemical principles. Boltz-2 yielded similar results (see SI Figure S4).

Further, we developed DexDesign (1, 24), which is a physics-based algorithm specialized for designing D-peptide competitive inhibitors. This algorithm can predict D-peptide:L-protein complexes and is guaranteed to have a 0% chirality violation rate. DexDesign exhibits structural recovery of known D-peptide binders and predicts low-energy conformations for drug complexes; these metrics for analyzing D-peptide ligands are reported in a restitution/replication framework (see Discussion section of DexDesign (1)). DexDesign therefore reports on the fold and binding pose of D-peptide ligands: the algorithm not only recovers known D-space chemical interactions but also returns metrics on the quality of D-peptide binders based on energy. Therefore, DexDesign is a viable methodology to complement AF3 by returning readouts of D-peptides, characterizing the binding pose and fold based on chemical feasibility. DexDesign is a step toward robust heterochiral design by providing indirect, physics-based readouts of structure quality. Further, DexDesign supplements AF3 with additional capabilities: this algorithm is capable of evaluating structure-disrupting mutations, which is an inherently difficult problem for deep learning models (9). DexDesign and subsequent extensions of the protocol (25) also provide structural ensemble outputs ordered by decreasing binding affinity.

To address low correlation for confidence metrics, future AlphaFold models should include metrics that explicitly assess chirality in predictions. Despite chirality violation incurring the highest penalty during benchmarks (8), the AF3 ranking score does not include a component that accounts for chirality: at no point after inference does AF3 explicitly confirm that input chirality specifications are respected in the output. This can be resolved by including a component in the ranking score that strongly penalizes predictions with incorrect stereocenters. In this way, chiral molecules with incorrect stereocenters will accurately receive lower ranking scores. This will improve post hoc assessment of chirality violations by AF3.

## Discussion

In this study, we systematically test the ability of AF3 to handle the chirality of ligands. Despite claims in the original paper (8) that AF3 is capable of handling diverse chemical interactions, we report the failure of this architecture on D-peptides, which has profound implications for predictions of drug complexes. Prior to this publication, the large interdisciplinary protein design community had only one metric for chirality violation: the 4.4% originally reported (8) in 2024. This work provides additional metrics on D-peptide ligand predictions that highlight the limitations of AF3 and proposes solutions using diffusion steering and physics-based algorithms. It appears AF3 has learned features that do not generate D-peptides with correct chirality.

While the previously reported (8) AF3 chirality violation rate was low (4.4%) across all chiral centers, we have discovered that the model fails to maintain chiral centers specified in the input for D-peptides. The average chirality violation rate across all samples is 51%, so the model is as accurate on average as chance (random chirality choice, L or D, for each peptide residue). Additionally, AF3 inaccurately folds and docks D-peptides in complex with L-targets. As shown in Figures 2 - 4, AF3 typically either places the D-peptide in the incorrect binding position or improperly orients the D-peptide in the correct binding interface. For the three D-peptides evaluated, each was folded incorrectly and was not predicted in the correct binding pose in any of the 15 samples (see SI Figure S1). The top-ranked DP19 sample was oriented upside down and on the opposite side of L-19437 (Figure 2). The top-ranked DP9 sample was rotated in the binding pocket such that hydrophobic residues were unfavorably solvent exposed (Figure 3). Further, DP12 sample 3’s backbone was rotated such that only three out of seven chemical contacts with MDM2 were recapitulated (Figure 4).

Interestingly, removing (apo) or changing the D-peptide’s target (GB1 or Ubiquitin, see SI Table S2) fails to reduce average chirality violation rate to anything approaching the 4.4% originally reported (8). It is possible that AF3’s L-residue training data led the model to conflate a biological principle—that proteinogenic amino acid stereocenters are L— with a false physical constraint that amino acid stereocenters must be L. Additionally, describing D-stereocenters with fluorines in place of *α*-hydrogens did not eliminate chirality violation in any tested system (SI Table S2). Thus, AF3 architecture may have difficulty maintaining stereocenters for organic molecules resembling D-peptides. These results indicate AF3 struggles with D-peptide chirality regardless of target, even when hydrogens are replaced with other small atoms.

Analysis of confidence metrics revealed no statistical correlation between chirality violation rate and confidence scoring (Figure 6). Further, although the fold and binding pose of each D-peptide sample fail to converge in a structural ensemble (see SI Figure S1), each D-peptide seed converges on low-confidence scoring (Figure 6 (C)). Because of the lack of correlation between D-peptide sample confidence rank and chirality violation rate, AF3 is unsuitable for D-peptides because its predictions are not only incorrect in chirality, fold, and pose, but also converge to low-confidence scoring. Unfortunately, these errors do not resolve with more seeds; as presented in Figure 5, the top-ranked and average chirality violation rates do not improve with additional predictions. This indicates the computational failures for this class of molecule cannot be resolved with larger prediction volumes.

While AlphaFold has enabled a wide range of research for varying design objectives (26–28), the model is currently unable to reliably predict D-peptides. These shortcomings may be attributed to D-peptide crystal structure scarcity or inherent model flaws. AF3 correctly predicts low confidence ranges for incorrect D-peptide structures, but it nevertheless fails to properly rank and predict the correct chirality, fold, and binding pose of D-peptide predictions, regardless of the number of predictions generated. This inability to produce high-quality D-peptide predictions indicates that further work is needed to effectively predict heterochiral complexes. It is possible that performance will improve with comprehensive chirality violation parameters and more empirical D-peptide:L-protein structural data to be used in future model training.

## Materials and Methods

The model parameters and AF3 inference pipeline were obtained from Google DeepMind (29) and run via a local implementation. Boltz-1x was obtained from GitHub (30) and run via a local implementation. Evaluated D-peptide:L-protein systems were obtained from the Protein Data Bank (12). These included DP19:L-19437 (PDB ID 7YH8), DP9:Streptavidin (PDB ID 5N8T), and DP12:MDM2 (PDB ID 3IWY). Water molecules and duplicate chains were removed. The D-peptide and L-target were split and saved into separate PDB files. Open Babel (31) was used to obtain the sequence of the L-protein and D-peptide. The sequence for SH3 and decoy L-targets GB1 and Ubiquitin were obtained with the same methodology. The D-peptide amino acid sequence was converted to Simplified Molecular Input Line Entry System (SMILES) data format (as used for AlphaFold 3 baseline models (8)) using NovoPro PepSMI (32). To ensure D-peptide SMILES data was correctly converted, chiral centers were confirmed by two methods: (1) confirmed in 2D using RCSB PDB Chemical Sketch Tool (33) and (2) confirmed in 3D by converting SMILES to PDB using NovoPro smiles2pdb (34) and visualizing in PyMOL (35). For D-peptides with fluorines replacing *α*-hydrogens, SMILES strings were manually revised to represent this substitution. These data inputs were similarly evaluated in 2D and 3D. The L-target sequence and D-peptide SMILES string were provided in the required format: AF3 in JSON, Boltz-1x in FASTA, and Boltz-2 in YAML. All Boltz-1x and Boltz-2 predictions were obtained using inference-time potentials. Unless otherwise specified, each job was run with one seed. Outputs were protonated with Protoss (15, 16) and inspected for L, D, and planar chiral centers.

## Data Availability

The Protein Data Bank (PDB) files used in this work can be accessed through the RCSB PDB (https://www.rcsb.org/) using the IDs referenced in the manuscript.

## ACKNOWLEDGMENTS

We received funding from the NIH (grant R35 GM-144042 to BRD; grants R01 GM-115355 and R01 AI-094475 to PZ). We thank Allen McBride, Liyang Zhu, Edward Cheng, Ying Sun, Edward Moseley, Eric Chen, and Yuxi Long for feedback on earlier versions of this manuscript.

## Supplementary Information

### 1. Decoy Targets and Fluorinated Peptides

In addition to testing each D-peptide in complex with the endogenous target and testing apo D-SH3, we placed each D-peptide in complex with decoy target Ubiquitin (PDB ID 6NXL, chain A) or GB1 (PDB ID 2J52, chain A) to further investigate if target affects ligand stereochemistry in AF3. Although predicting D-peptides with endogenous target and apo failed to improve chirality violation rate to anything approaching 4.4%, we evaluated if non-endogenous L-targets would improve predictions. While incorrect ligand chirality for an AF3 D-peptide:L-endogenous target may indicate a faulty chirality interpretation module, an incorrect D-peptide:L-decoy provides further evidence that AF3 has a fundamental flaw with D-peptides. We report that AF3 predicts models with high chirality violation (average 44% across all decoy systems) and again displays inconsistent patterns of D/L residue chirality for a given D-peptide. Additionally, to test whether AF3 chirality violation was due to hydrogen stripping in AF3’s input processing, we replaced each D-peptide *α*-hydrogen with a different small atom, fluorine. When placed in complex with the respective endogenous target, AF3 still produces models with a high chirality violation (average 33% across all three systems) despite non-hydrogen stereocenters. These results are displayed in SI Table S2.

**Table S1.**
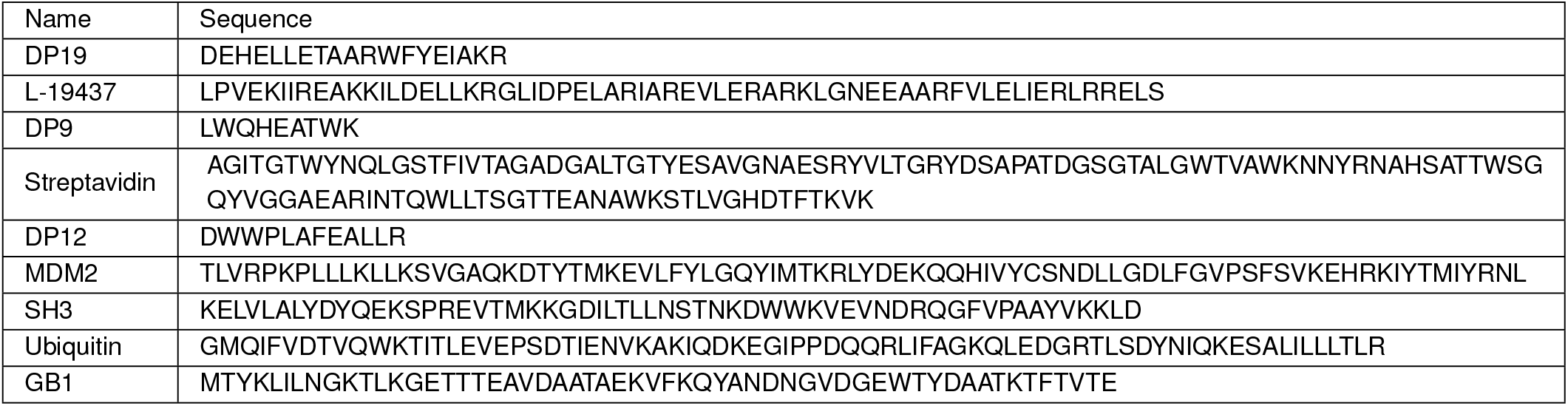
Full sequences for DP19:L-19437 (PDB ID 7YH8), DP9:Streptavidin (PDB ID 5N8T), DP12:MDM2 (PDB ID 3IWY), SH3 (PDB ID 1SHG), and decoy targets Ubiquitin (PDB ID 6NXL) and GB1 (PDB ID 2J52). The Name column indicates protein label and the Sequence column contains the sequence in single-letter amino acid notation.

**Table S2.**
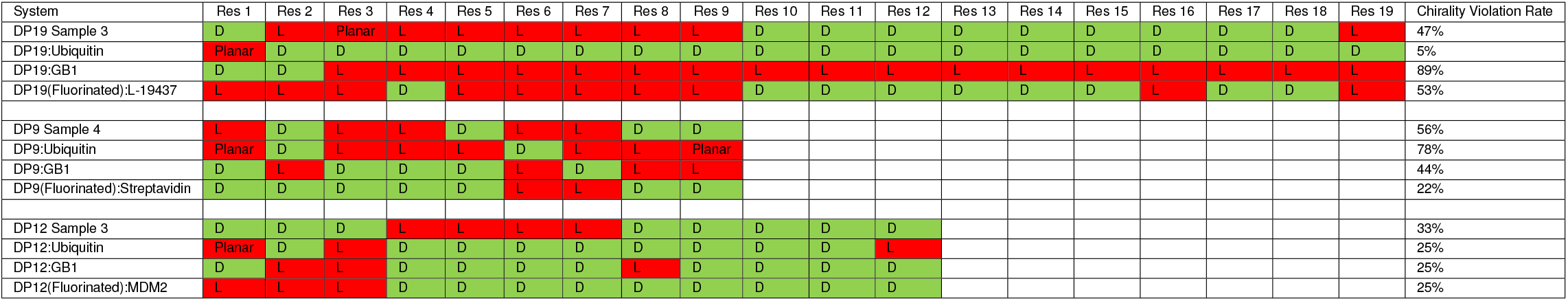
Per-residue chirality violations for AlphaFold 3 predictions of D-peptides with synthetic L-targets or fluorine. The System column details the evaluated complex. DPX:Ubiquitin is the D-peptide DPX (DP19, DP9, or DP12) in complex with Ubiquitin. DPX:GB1 is the D-peptide DPX (DP19, DP9, or DP12) in complex with GB1. Systems with “(Fluorinated)” indicate the D-peptide’s *α*-hydrogens were replaced with fluorine. Top-ranked samples from Table 1 are repeated here for reference. For brevity, only the top-ranked sample for each experiment is displayed. The Res # columns indicate the chirality of the residue index (D (correct) or L (incorrect)). Entries of “Planar” indicate that AlphaFold 3 incorrectly predicted a flattened protein backbone that could not be assigned to either chirality (see Figure 1). Correct D stereocenters are in green while incorrect L and Planar stereocenters are in red. Chirality violation is calculated by dividing the number of peptide residues with incorrect chirality (L or Planar) by the length of the peptide. Decoy systems (peptides targeting GB1 or Ubiquitin) display inconsistent patterns of D/L chirality violation, per residue, on the top-ranked model. Describing chiral centers with fluorines in place of *α*-hydrogens failed to eliminate chirality violation in any system.

**Fig. S1.**
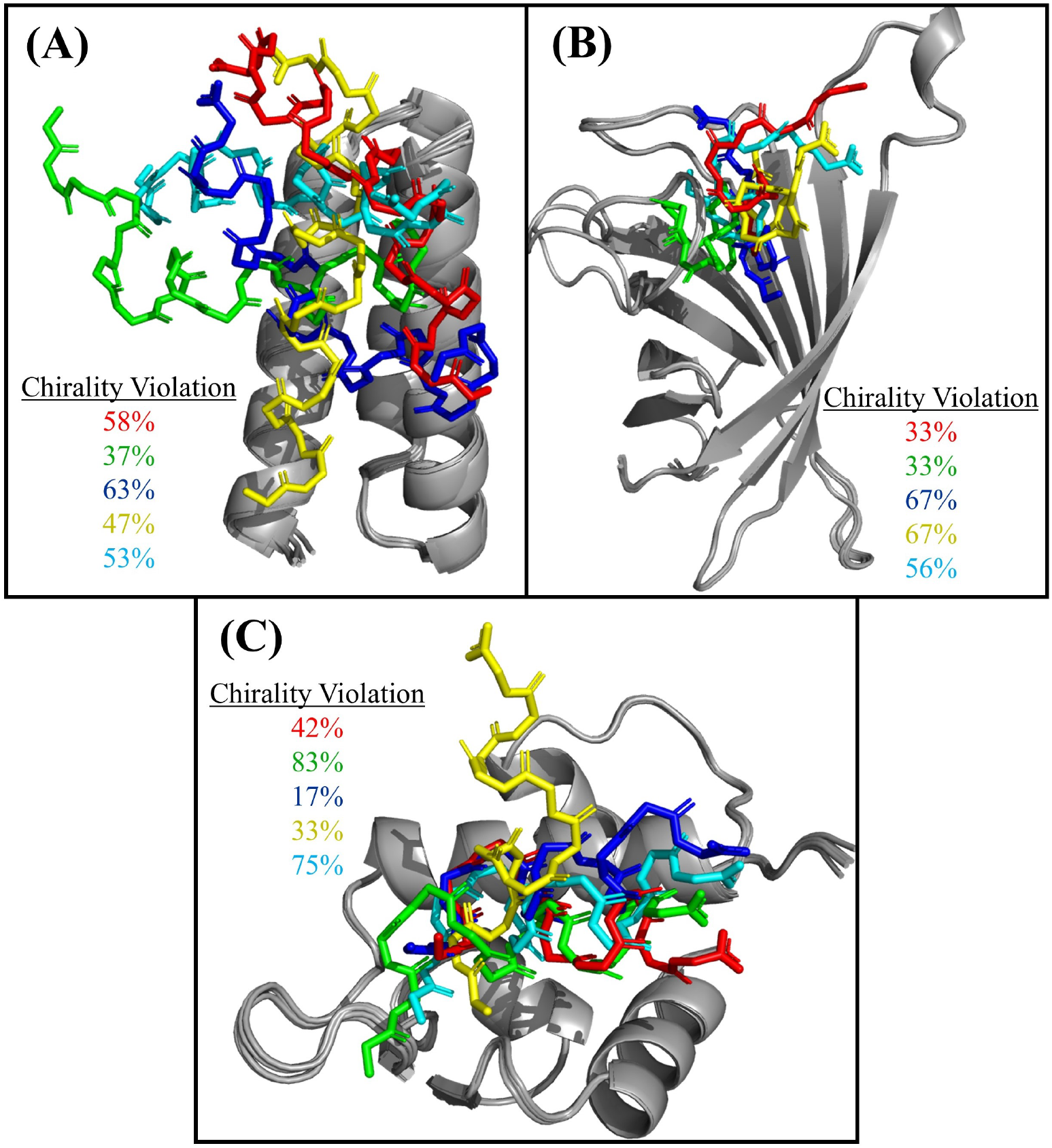
Samples from each AlphaFold 3 D-peptide prediction fail to converge. **(A)**: AF3 predictions for DP19:L-19437 (PDB ID 7YH8). **(B)**: AF3 predictions for DP9:Streptavidin (PDB ID 5N8T). **(C)**: AF3 predictions for DP12:MDM2 (PDB ID 3IWY). Each frame includes the five samples produced by AF3 (see Table 1 for chirality violation rates). All L-targets are shown as gray cartoons, while D-peptide samples are shown as sticks. Sidechains are hidden for clarity. Sample 0 is red, sample 1 is green, sample 2 is blue, sample 3 is yellow, and sample 4 is cyan. Chirality violation rates for the D-peptides are reported in each frame as the corresponding sample color. The L-target predictions largely converge; the set of samples for each system demonstrates few structural deviations. While AF3 docks each ligand prediction sample in the same general binding site, the D-peptide samples in each system fail to achieve a consensus fold or binding pose. (A): predictions for DP19 deviate in structure significantly between each sample. In addition to docking the predictions in the incorrect binding location (see Figure 2 for more details), each sample is predicted in a different orientation in the binding pocket, resulting in a C-term distance between sample 0 and sample 1 of 32.7 Å. (B): similar to DP19, predictions for DP9 fail to reach a fold and pose consensus (see Figure 3 for more details). The C-term distance between sample 0 and sample 1 is 14.5 Å. (C): DP12 fails to converge in fold or binding pose (see Figure 4 for more details). The C-term distance between sample 0 and sample 3 is 18.4 Å.

**Fig. S2.**
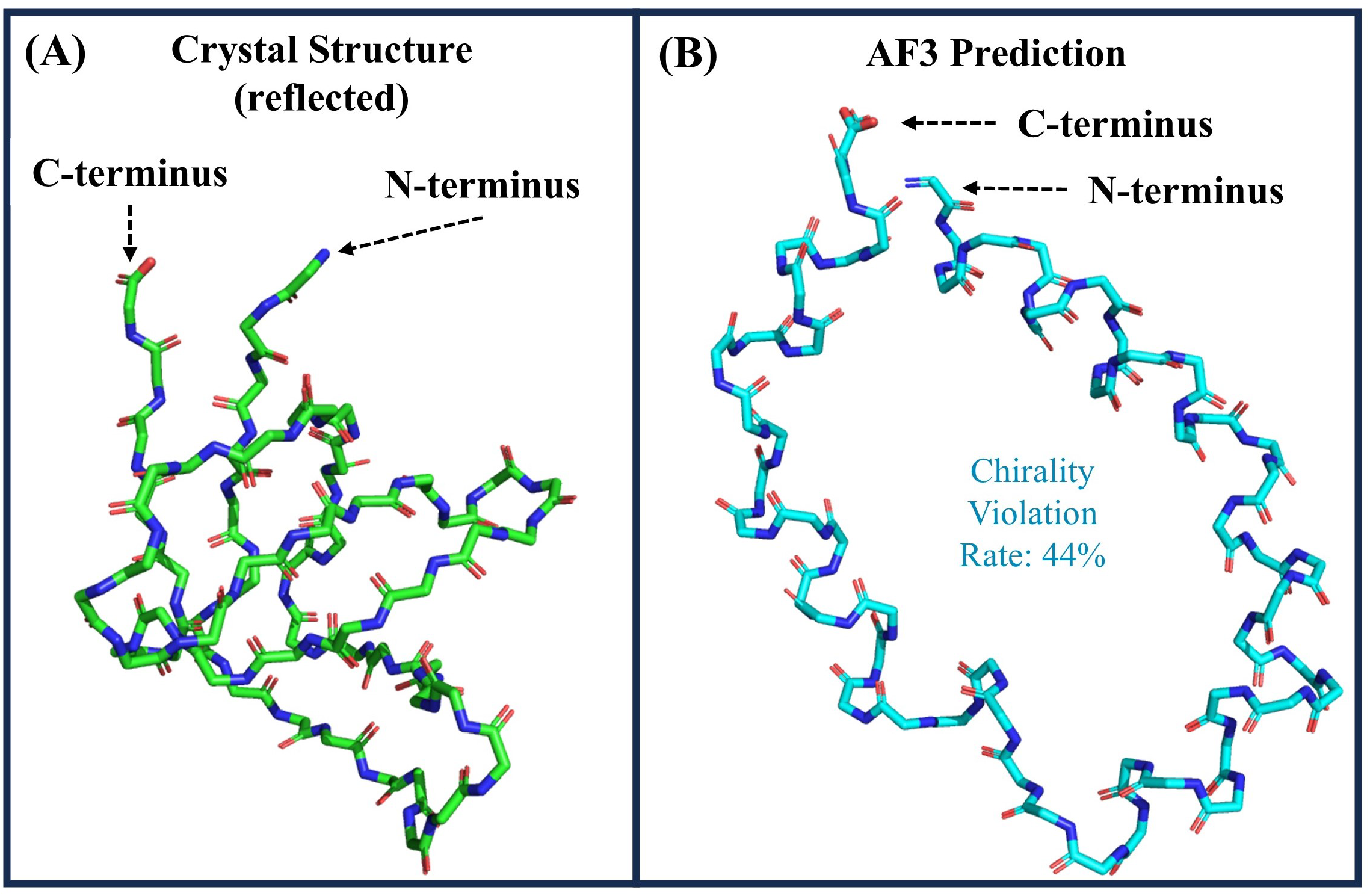
Comparison of D-SH3 crystal structure (PDB ID 1SHG, reflected across the z-axis) to AF3 top-ranked (sample 0) prediction of D-SH3. Sidechains are hidden for clarity. **(A)**: The crystal structure of Src-homology 3 (SH3) reflected into D-space (D-SH3, green sticks). L-SH3 and D-SH3 should adopt the same fold, but with opposite handedness (1, 2). **(B)**: AF3 prediction of D-SH3 (cyan sticks). This structure differs significantly from known crystal structure (C*α* alignment RMSD of D-SH3 to AF3 prediction is 13.9 Å) and has a chirality violation rate of 44%.

**Table S3.** Confidence metrics returned by AlphaFold 3 for predictions of DP19:L-19437 (PDB ID 7YH8), DP9:Streptavidin (PDB ID 5N8T), and DP12:MDM2 (PDB ID 3IWY). The System column indicates the evaluated complex and sample number. Ranking score is calculated by AF3 using the formula 0.8(ipTM) + 0.2(pTM) + 0.5(fraction_disordered) −100(has_clash). The D-peptide pTM, L-protein pTM, and Complex pTM columns indicate the predicted template modeling (pTM) score for the peptide, protein, and complex, respectively. Derived from template modeling (TM) (3) score, the pTM score reports how confident AF3 is in the accuracy of the entire structure. A pTM score ≥0.5 means the overall structure is likely similar to the crystal structure (4). The D-peptide ipTM, L-protein ipTM, and Complex ipTM report the interface predicted template modeling (ipTM) score for the peptide, protein, and complex, respectively. The ipTM score reports the confidence in predicted interfaces. Because AF3 is only evaluating a single D-chain:L-chain interface, these values will be identical for a given sample. ipTM scores *>* 0.8 are considered high-quality predictions, values from 0.6−0.8 are a “grey zone” where accuracy is uncertain, and values ≤0.6 likely indicate a poor prediction (4). The fraction_disordered column reports a scalar in the range 0 −1 that indicates the fraction of prediction structure that is disordered. The has_clash column is a boolean indicating if the predicted sample contains either a chain with 100 clashing atoms or a chain where *>* 50% of the chain is clashing. AF3 correctly predicts high confidence for the L-target samples. AF3 correctly reports low confidence ranges for each D-peptide sample, but incorrectly ranks individual samples according to chirality violation rate.

| System | Ranking Score | D-Peptide Chain pTM | L-Protein Chain pTM | Complex pTM | D-peptide Chain ipTM | L-Protein Chain ipTM | Complex ipTM | fraction_disordered | has_clash |
| --- | --- | --- | --- | --- | --- | --- | --- | --- | --- |
| 7YH8 Sample 0 | 0.61 | 0 | 0.80 | 0.68 | 0.59 | 0.59 | 0.59 | 0 | 0 |
| 7YH8 Sample 1 | 0.74 | 0 | 0.83 | 0.79 | 0.73 | 0.73 | 0.73 | 0 | 0 |
| 7YH8 Sample 2 | 0.53 | 0 | 0.77 | 0.62 | 0.50 | 0.50 | 0.50 | 0 | 0 |
| 7YH8 Sample 3 | 0.77 | 0 | 0.79 | 0.81 | 0.76 | 0.76 | 0.76 | 0 | 0 |
| 7YH8 Sample 4 | 0.67 | 0 | 0.79 | 0.73 | 0.66 | 0.66 | 0.66 | 0 | 0 |
| 5N8T Sample 0 | 0.60 | 0.42 | 0.88 | 0.75 | 0.56 | 0.56 | 0.56 | 0 | 0 |
| 5N8T Sample 1 | 0.66 | 0.47 | 0.87 | 0.78 | 0.63 | 0.63 | 0.63 | 0 | 0 |
| 5N8T Sample 2 | 0.54 | 0.33 | 0.87 | 0.72 | 0.49 | 0.49 | 0.49 | 0 | 0 |
| 5N8T Sample 3 | 0.57 | 0.45 | 0.87 | 0.74 | 0.53 | 0.53 | 0.53 | 0 | 0 |
| 5N8T Sample 4 | 0.68 | 0.45 | 0.87 | 0.79 | 0.65 | 0.65 | 0.65 | 0 | 0 |
| 3IWY Sample 0 | 0.62 | 0 | 0.82 | 0.72 | 0.59 | 0.59 | 0.59 | 0 | 0 |
| 3IWY Sample 1 | 0.61 | 0 | 0.81 | 0.72 | 0.59 | 0.59 | 0.59 | 0 | 0 |
| 3IWY Sample 2 | 0.59 | 0 | 0.81 | 0.70 | 0.57 | 0.57 | 0.57 | 0 | 0 |
| 3IWY Sample 3 | 0.66 | 0 | 0.83 | 0.75 | 0.64 | 0.64 | 0.64 | 0 | 0 |
| 3IWY Sample 4 | 0.62 | 0 | 0.82 | 0.73 | 0.60 | 0.60 | 0.60 | 0 | 0 |

**Fig. S3.**
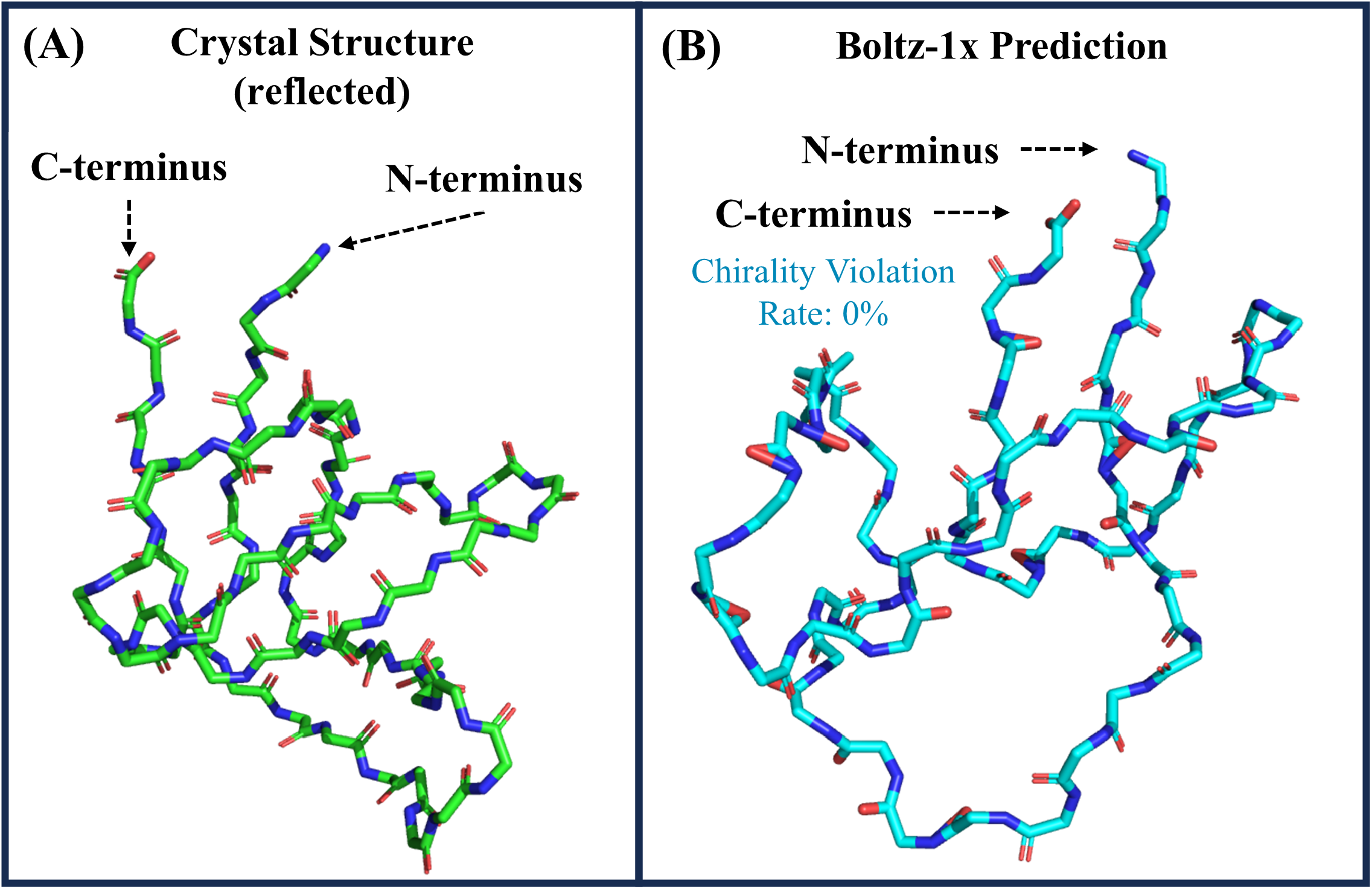
Comparison of D-SH3 crystal structure (PDB ID 1SHG, reflected across the z-axis) to Boltz-1x top-ranked prediction of D-SH3. Sidechains are hidden for clarity. **(A)**: The crystal structure of Src-homology 3 (SH3) reflected into D-space (D-SH3, green sticks). L-SH3 and D-SH3 should adopt the same fold, but with opposite handedness (1, 2). **(B)**: Boltz-1x prediction of D-SH3 (cyan sticks). The C*α* alignment RMSD of D-SH3 to Boltz-1x prediction is 13.3 Å. Although the fold is similarly inaccurate as AF3 (see SI Figure S2), the chirality violation rate across all 57 residues is 0%. This is likely due to the fact that Boltz-1x uses a chemistry-inspired potential function (5) that tilts the distribution to respect chemical principles.

**Fig. S4.**
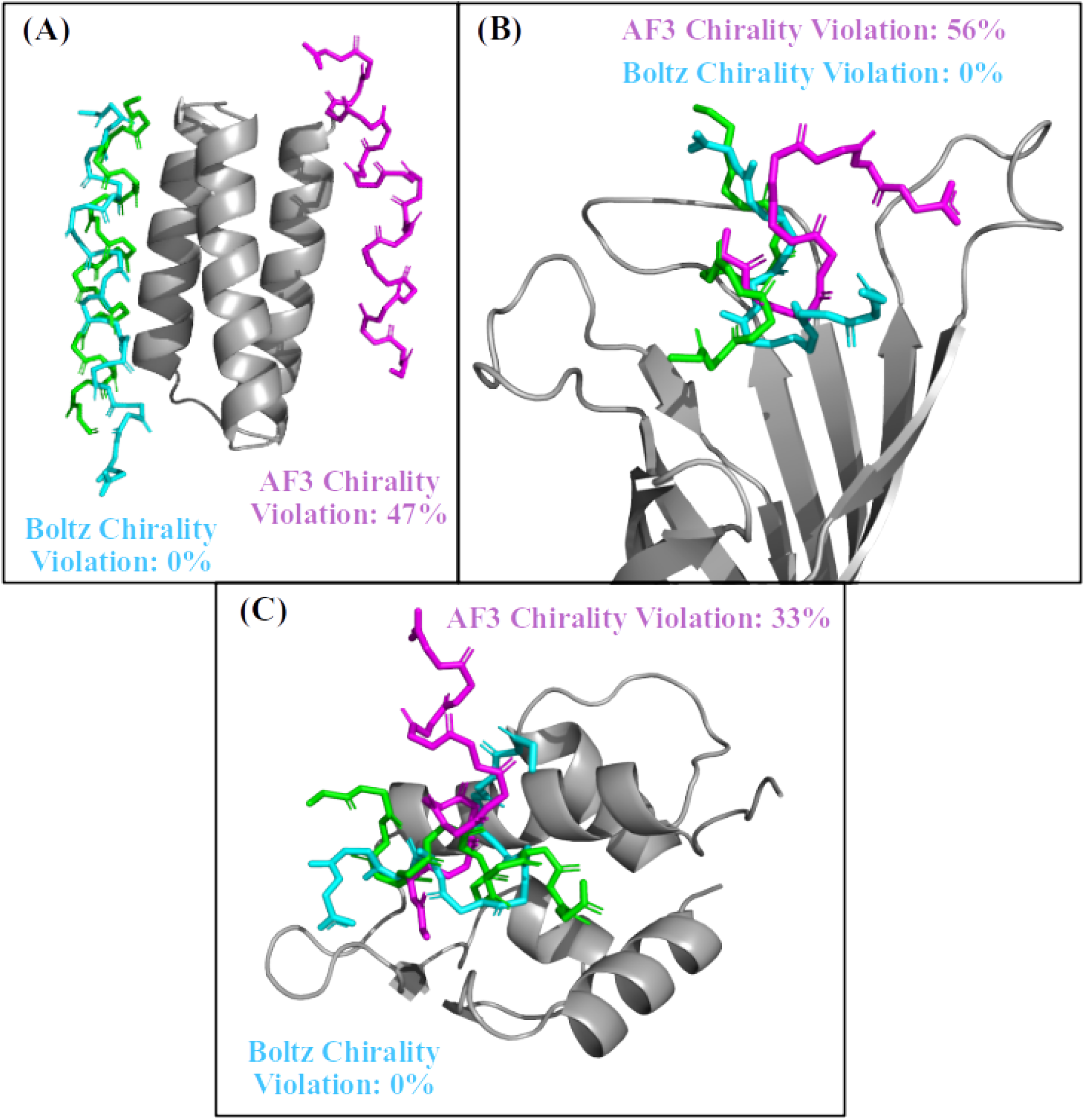
Comparison of Boltz-2 (with inference-time potentials) and AF3 top-ranked predictions to crystal structures for D-peptide:L-protein complexes. **(A)**: Comparisons for DP19:L-19437 (PDB ID 7YH8). **(B)**: Comparisons for DP9:Streptavidin (PDB ID 5N8T). **(C)**: Comparisons for DP12:MDM2 (PDB ID 3IWY). The L-target in each frame is shown in grey cartoon, and the D-peptides are in stick representation. The crystal structure is in green, Boltz-2 is in cyan, and AF3 is in violet. Sidechains are hidden for clarity. With the exception of DP19, Boltz-2 predictions of the D-peptides are about as accurate as Boltz-1x with respect to fold and binding pose (see Figure 7). The chirality violation rate of Boltz-2 (with inference-time potentials) for all systems was 0%.

